# Morphogenesis of the Carapace from Phyllosoma to Puerulus in Spiny Lobsters

**DOI:** 10.1101/2025.11.20.688360

**Authors:** Haruhiko Adachi, Kentaro Morikawa, Yasuhiro Inoue, Shigeru Kondo

**Affiliations:** Institute for Advanced Biosciences, Keio University, Tsuruoka, Yamagata 997-0017, Japan; Graduate School of Media and Governance, Keio University, Fujisawa, Kanagawa, 252-0882, Japan; Department of Micro Engineering, Graduate School of Engineering, Kyoto University, Kyoto, Kyoto 616-8540, Japan; National Institute of Genetics, Mishima, Shizuoka, 411-8540, Japan

**Keywords:** Phyllosoma, Puerulus, Metamorphosis, Morphogenesis, Molting

## Abstract

Molting in exoskeleton-bearing organisms can result in dramatic body shape transformations. While this phenomenon has been well-studied in insects, limited knowledge exists for other taxa, including crustaceans. Among crustaceans, achelatan lobsters, such as Palinuridae and Scyllaridae, exhibit a unique metamorphic transition from a flattened phyllosoma larva to a three-dimensional puerulus or nisto larva in a single molt. This study investigates the mechanisms underlying this transformation, focusing on carapace deformation through histological analysis, live imaging, and computational modeling. Three-dimensional micro-computed tomography revealed a significant reduction in body length during metamorphosis. Live imaging captured dynamic morphological changes, including epithelial contraction and subsequent reshaping. Scanning electron microscopy identified a distinctive furrow structure on the carapace surface, absent in post-molt puerulus larvae, suggesting a role in the transformation process. To explore the formation of this furrow pattern, a computational model was developed based on differential shrinkage rates in vertical and horizontal directions. The model successfully reproduced similar patterns observed in natural specimens, implying that controlled anisotropic contraction could contribute to morphogenesis. Furthermore, three-dimensional shrinkage simulations demonstrated that local contraction with constrained out-of-plane deformation can generate folds, which later expand to form curved structures. This study provides novel insights into the biomechanics of arthropod molting, highlighting a previously unrecognized mechanism of two-dimensional-to-three-dimensional transformation. The findings enhance our understanding of achelatan lobster development and offer broader implications for exoskeletal morphogenesis across arthropods.

## Introduction

In some exoskeletons, molting can cause extreme changes in body shape. The mechanism has been studied in detail in insects, including flies(de la Loza and Thompson 2017) and beetles(Matsuda, Gotoh et al. 2017, Gotoh, Adachi et al. 2021), but little is known about other species. Crustaceans also exhibit diverse larval morphologies depending on the species and undergo drastic morphological changes through the process of molting (Møller, Anger et al. 2020). Among crustaceans, achelata have the characteristic larval style. Achelata includes Palinuridae and Scyllaridae, both of which undergo a very flat morphology (<u>Mov. 1</u>), called phyllosoma larvae, although their outlines differ (Landeira, Deville et al. 2023). After repeated molting, the final instar phyllosoma larvae undergo a metamorphosis in which they transform into puerulus larvae in Palinuridae and nisto larvae in Scyllaridae with a single molt, and then grow through juvenile shrimp (Matsuda, Takenouchi et al. 2006, Wakabayashi and Phillips 2016). In other words, the process of transformation from a flat, two-dimensional form to a three-dimensional form occurs in a single molting. In order to elucidate the mechanism of transformation from a flat to a three-dimensional state, we focused our attention on the morphogenesis of this spiny lobster.

With regard to the phyllosoma to puerulus metamorphosis, temporal transcriptome analysis *Sagmariasus verreauxi* and *Panulirus ornatus* has been conducted, and sequences that are characteristically expressed at each stage have been found (Ventura, Fitzgibbon et al. 2015, Hyde, Fitzgibbon et al. 2019). The cross-sectional structure of the cuticle at each stage has also been analyzed, and the oriented chitin fiber structure seen in general crustaceans has not been observed in phyllosoma larvae, and a characteristic amorphous calcium carbonate structure has been confirmed. Then, although calcium carbonate usually has a crystalline structure, proteins that are thought to be important for maintaining an amorphous state in phyllosoma have also been identified (Ventura, Nguyen et al. 2019). In addition, changes in the elemental composition during the development of phyllosoma have recently been captured (McDougall, Deas et al. 2023). Aquaculture studies have also been conducted in Japan for a long time in species such as *Panulirus japonicus* and *Panulirus penicillatus*, and the developmental process including the larval stage has been described (Kittaka and Kimura 1989, Matsuda, Takenouchi et al. 2006, Matsuda and Takenouchi 2007), as well as the observation of the dynamics during molt (Murakami, Jinbo et al. 2007). On the other hand, the curvature of the carapace, which is characteristic of the transformation from phyllosoma to puerulus, remains poorly understood. In this study, we focused on the deformation of the carapace, analyzed what kind of deformation is occurring using histological analysis and live observation at the time of molting, and proposed a model of the deformation.

## Materials and Methods

### Animals

*Panulirus japonicus* lobsters were cultured at the Mie Prefectural Science and Technology Promotion Center, Fisheries Research Division. The culture conditions were identical to those employed in previous studies (Matsuda and Takenouchi 2007). The specimens were fixed in 70% ethanol. Molting was recorded using two video cameras: an FDR-AX (Sony, Japan) and an iPhone SE2 (Apple, USA). Molting onset was determined by antenna folding, following a previous study (Murakami, Jinbo et al. 2007).

### Micro-CT (µCT)

Following established methods in insect studies (Adachi, Matsuda et al. 2020), specimens fixed in 70% ethanol were transferred to 100% ethanol and finally to t-BtOH. Specimens were then frozen at 4°C and dried using a freeze dryer to minimize deformation. Dried specimens were scanned using a µCT Skyscan 1172 (Bruker, Belgium) and reconstructed with NRecon ver. 1.7.0. Mesh files were also output as stl data for later 3D simulations. Cross-sectional images were obtained using CTAn, and length measurements were taken with Fiji (Schindelin, Arganda-Carreras et al. 2012).

### Scanning Electron Microscopy (SEM)

CT-scanned specimens were sputter-coated with gold and used for SEM analysis. They were attached to carbon seals and imaged at 10 kV on a JCM-6000 NeoScope (JEOL, Japan). The resulting images were concatenated using the Fiji plugin Image Stitching (Preibisch, Saalfeld et al. 2009).

### Expansion Simulation

This followed methods established for beetle horns (Matsuda, Gotoh et al. 2017, Matsuda, Gotoh et al. 2021). Calculations were performed to flatten the angle between adjacent triangular meshes while preserving their area. Implementation was carried out in Julia language ver 1.8.

### Two-Dimensional Computational Shrinkage

It was performed in Julia language ver 1.8. Displacements of ellipses and rectangles were placed on each side of the rectangle and distributed inside the rectangle to simulate a quasi-contraction (Fig. S2). The distribution was carried out according to a gaussian distribution, in which the internal influence is varied by the variance parameter σ. Protruding ends were distributed vertically and horizontally based on the aspect ratio during the initial arrangement and internally according to a Gaussian distribution. When σ is small, the pattern stops on the porch, and when σ is large, it penetrates more inside.

### Three-Dimensional Computational Shrinkage

We represented the carapace epithelium as a triangular mesh and simulated its deformation by computing the motion of mesh vertices based on a mechanical model. The motion of each vertex was assumed to obey an overdamped (viscous-dominated) equation of motion, and the force acting on each vertex was given by the gradient of the total energy, which consisted of stretching, bending, and *z*-directional displacement-constraint energy terms (Inoue, Tateo et al., 2020). The *z*-directional displacement-constraint energy represents the restriction of out-of-plane deformation imposed by the cuticle. This model was used both for simulations of furrow formation induced by shrinkage and for simulations of furrow unfolding.

### Equation of motion for vertex *i* of a triangular mesh

Assuming viscous-dominated dynamics, the equation of motion for vertex *i* with position ***x***_*i*_ is written as

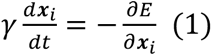

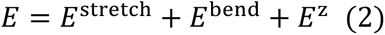

where 𝛾 is a friction coefficient and *E* is the total energy. In all simulations, we set 𝛾 = 1.0.

### Details of each energy function

#### Energy of stretching (Chen, Sastry et al. 2018)

The in-plane strain 𝜺_*i*_ due to stretching in the triangular mesh was expressed, for each triangle *i*, using its discrete first fundamental form ***I***_*i*_ as follows:

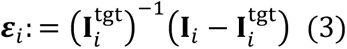

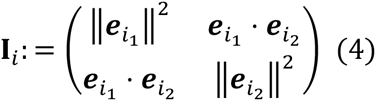

where 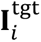 is the first fundamental form of triangle *i* in the reference (target) configuration and 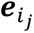 denotes the vector of edge *j* of triangle *i*. The stretching energy was written using material parameters 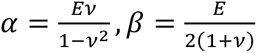 where ***E*** is the Young’s modulus and 𝜈 is the Poisson’s ratio:

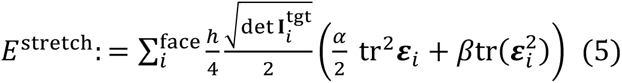

where ℎ denotes the thickness. In all simulations, we set 𝛼 = 6.6 × 10^3^, 𝛽 = 7.7 × 10^3^, ℎ = 1.0 × 10^-2(^. In the shrinkage simulations, to realize anisotropic shrinkage, we set 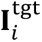 as the shape obtained from the initial shape 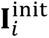 by scaling it by a factor 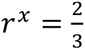 in the 𝑥-direction (the major axis of the initial elliptical shape) and by a factor 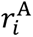 in area. The target area was thus written as 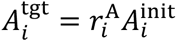; the procedure for specifying 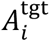 is described below.

#### Energy of bending (Tamstorf and Grinspun 2013)

As the bending energy of the triangular mesh, we adopted an energy function based on the dihedral angle 𝜃_*i*_ between the two triangles sharing each edge *i*:

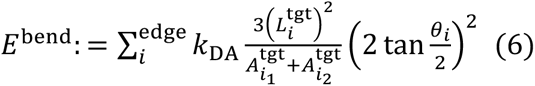

where 𝑘_DA_ is an elastic coefficient, 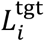^"^ is the length of the edge *i* in the reference configuration, and 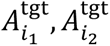 are the areas of triangles sharing the edge *i* in the reference configuration. In the shrinkage simulations, this term ***E***^bend^ was interpreted as the mechanical bending energy of the material. In the unfolding simulations, the same functional form was used as a geometric flattening energy to obtain the unfolded shape. We set 𝑘_DA_ = 1.0 × 10^-4^ in the shrinkage simulations and 𝑘_DA_ = 1.0 × 10^-2(^ in the unfolding simulations.

#### Energy of 𝒛-directional displacement constraint (Inoue, Tateo et al. 2020)

In the shrinkage simulations, the initial configuration of the tissue was taken to be an elliptical domain in the 𝑥𝑦-plane, and we assumed that the cuticle constrains deformation in the *z*-direction. Accordingly, the *z*-directional displacement-constraint energy was defined as

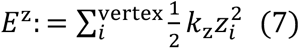

where *z*_*i*_ is the *z*-coordinate of vertex *i*. In the shrinkage simulations, we set 𝑘_z_ = 0.3. In the unfolding simulations, where the cuticle constraint is absent, we set 𝑘_z_ = 0.0.

#### Set-up formula for shrinkage rate distribution

In the shrinkage simulations, the target area 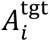 of each triangle *i*was defined relative to its initial area 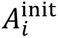 as

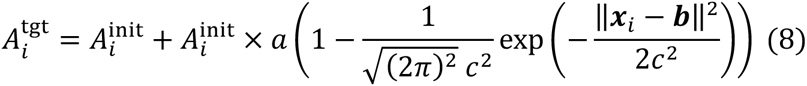

where ***x***_*i*_ is the position of triangle *i* (e.g. its centroid). This defines a shrinkage-rate distribution in which the target area reduction gradually increases with distance from the position 𝒃. The steepness of this gradient is controlled by the parameter 𝑐. In the simulations, we set 𝒃 = (1.1, 0.0) and 𝑐 = 0.5. The parameter 𝑎 determines the overall magnitude of shrinkage. Guided by experimental observations, we chose 𝑎 so that the total area changes in the simulations matched the experimentally observed total area change, i.e. such that 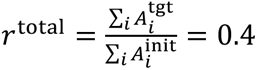.

## Results and Discussion

### Deformations During Metamorphosis

Firstly, in order to ascertain the nature of the occurring deformations, the shape of the final instar phyllosoma and puerulus larvae was examined (Fig. 1a). Stereomicroscopic images do not allow observation of deformation in the depth direction; therefore, the three-dimensional morphology was analyzed using μCT (Fig. 1b). By analyzing the length from the top of the head to the tail and the length divided at the mouthpart from the cross-sectional view of the midline, the lengths from the top of the head to the mouth, from the mouth to the tail, and the total length were determined (Fig. 1c). The length from the head to the tail decreased by approximately 73% from phyllosoma to puerulus (Fig. 1c). Similarly, the length from mouth to tail decreased by approximately 13% and the total length decreased by approximately 36%. Furthermore, the length of the dorsal side was analyzed in the lateral direction. This was done by analyzing cross-sectional views: the mouthpart in phyllosoma, and both the mouthpart and the posterior part of the carapace in puerulus—the widest area when viewed from above. As a result, lateral contraction was also observed. After contraction, the lengths at the mouthpart and posterior side were found to be approximately equal (Fig. 1c). Although previous studies have analyzed the morphology of phyllosoma and puerulus, the three-dimensional analysis using μCT focusing on the carapace represents a novel contribution to the field.

**Fig. 1.**
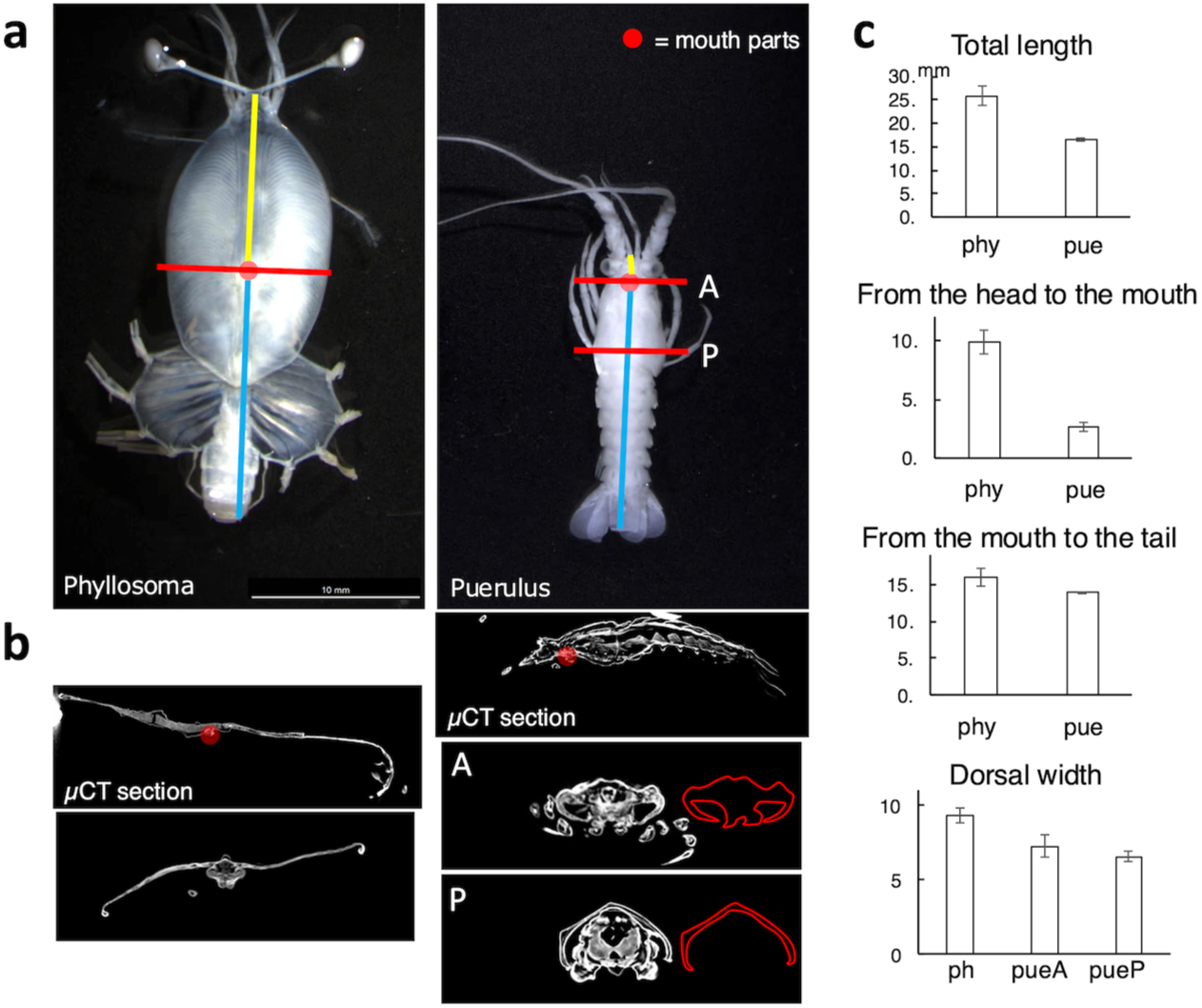
Morphology of phyllosoma and puerulus. (a) Brightfield images of phyllosoma and puerulus larvae. (b) Cross-sectional images using µCT, taken at the midline (yellow and blue lines in (a)), at the mouth organ (red points in (a)), and at the widest point of the carapace (red line P in (a)). (c) Quantification of phyllosoma and puerulus morphology. Measurements were taken of the ventral lateral body length at the midline cut surface, the length divided at the mouth organ, and the dorsal side of the transverse section (n=3). Although the sample size is small, the standard error is low, meaning we considered the results to be reliable.

In order to investigate the deformation on a shorter timescale, a video was taken at the time of molting (Fig. 2, <u>Mov. 2</u>). The entire process was successfully captured, beginning with the onset of the ocular peduncle falling and concluding with the puerulus form, which serves as a cue for the start of molting as previously observed in studies (Murakami, Jinbo et al. 2007). First, it was observed that the molting process, which lasts approximately 20 minutes, involves deformation from a planar structure to a shrimp-like form. Focusing on carapace deformation, we confirmed that the epithelial tissue first detached from the old cuticle and contracted (Fig. 2a,b, <u>Mov. 3</u>). It has been demonstrated by preceding studies that gut retracts require 3–4 days to reach completion (Ventura, Nguyen et al. 2019). On the basis of this timescale, it is therefore considered to be epithelial contraction. Subsequently, it was confirmed that the shrunken epithelial tissue was deformed (Fig. 2c). Then, we observed that the puerulus form developed following a pumping movement of the abdomen (Fig. 2d, <u>Mov. 4</u>). Previous studies have observed the movement of objects during molting (Murakami, Jinbo et al. 2007), but this study is the first to focus on the deformation of the carapace.

**Fig. 2.**
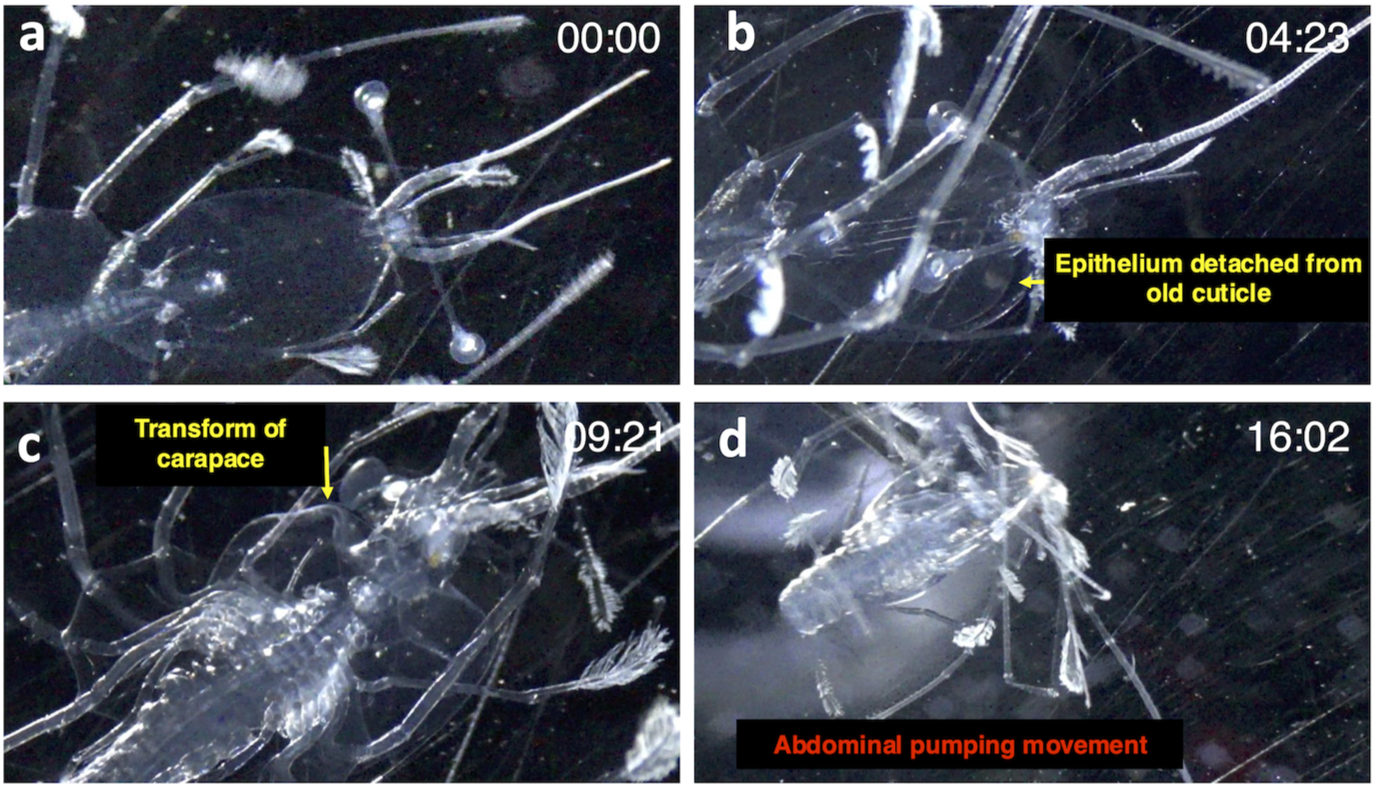
Live observation of the metamorphosis from phyllosoma to puerulus. (a) Early onset of metamorphosis. The ocelli, a signal for metamorphosis onset, are beginning to collapse. No significant changes are observed in the carapace. (b) 4 min 23 sec after (a). The carapace epithelium can be seen detaching from the old cuticle. (c) 9 min 21 sec after (a). Deformation of the carapace is observed. (d) 16 min 2 sec after (a). Pumping movements of the abdomen are observed.

To gain insight into the molting process, we conducted observations on individuals that had undergone the process but had not survived. The old cuticle was carefully removed through dissection, and the surface of the dorsal carapace was examined using scanning electron microscopy (SEM). This indicated a unique furrow structure, suggesting that it was formed during the molting process. (Fig. 3a). The dorsal carapace of puerulus larvae was also observed, but the above furrow structure was not confirmed (Fig. 3a). In insects, the folding and unfolding of the cuticle during molting has been the subject of study (Matsuda, Gotoh et al. 2017, Adachi, Matsuda et al. 2020). Recently, it has also been proposed that a comparable mechanism may be present in small freshwater shrimp (Adachi, Moritoki et al. 2025), thus suggesting that a similar phenomenon could occur during the transformation of phyllosoma to puerulus, which was the focus of this study. It is plausible that the pumping movement facilitates the unfolding of the furrow structure of the cuticle surface during molting. Indeed, a previous study by K. Matsuda et al. (2017) confirmed that furrow extension contributes to the final shape (Matsuda, Gotoh et al. 2017). Firstly, 3D mesh data was acquired by μCT for individuals in the process of molting whose furrow structure was confirmed, and the dihedral angle between the meshes was flattened. This revealed that the carapace was curved to some extent due to the extension of furrows (Fig. 3b). Conversely, this simulation also showed deformation in macroscopic structures that were not expected to extend, raising concerns about possible artifacts. Additionally, the computer simulations may use physical parameters that are unrealistic. To address these concerns, we used paper to reproduce the furrow pattern and perform an analog simulation of its extension. In the process of attempting to create a distinctive furrow pattern, it was determined that the upper portion would protrude. To address this issue, the protruding segment was secured to prevent movement. Additionally, it was observed that the original primordia does not exhibit furrows along the left and right edges. To address this, the edges were fixed to ensure they remained stationary. Consequently, even in the folding reproduced on paper, the curvature was reproduced to some extent by its unfolding (Fig. 3c). These results suggest that the characteristic folding observed in molting may contribute to the final shape. It is important to note that in previous studies of insects, it has been proposed that the expansion is due to the pressure of body fluid in the body cavity (Matsuda, Gotoh et al. 2017). This would suggest that the deformation is from three-dimension to three-dimension. However, this is not the case in this instance of phyllosoma. The deformation from two-dimension to three-dimension is a unique feature of the phyllosoma to puerulus deformation. In other words, the mechanism of extension of the folding structure is controversial. Furthermore, since phyllosoma is a planktonic organism that floats in the ocean, it is unclear how they achieve such a drastic deformation in the presence of the movement of the surrounding water. Additionally, puerulus larvae do not engage in feeding behavior for approximately 10 days until they molt into juvenile shrimp (Kittaka and Kimura 1989), which raises questions about the energy efficiency of the above transformation. In light of these intriguing questions, we hope that further analysis will be conducted.

**Fig. 3.**
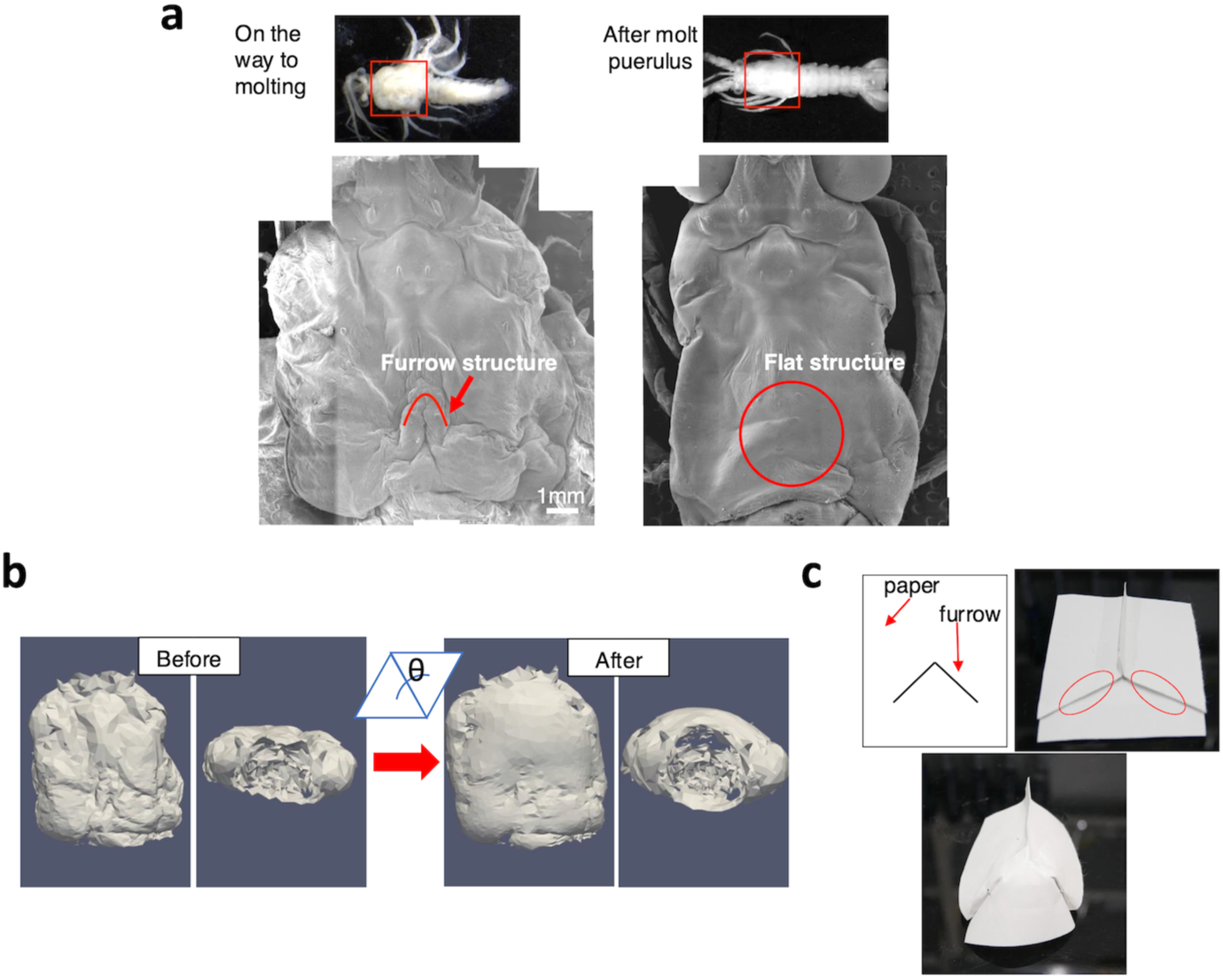
Transformation of the cuticular surface structure and furrow extension simulation (a) Cuticular surface of mid-molt and post-molt puerulus. Distinctive furrow structures can be observed on the mid-molt, but not on the post-molt individuals. (b) Computational furrow extension on the carapace of a mid-molt individual reconstructed using µCT. The entire carapace is curved due to furrow extension. (c) Reproduction of furrow patterns on paper and their unfolding process. By unfolding the reproduced furrow on paper, the curvature can be recreated.

### Consideration of how the furrow pattern forms

Next, we consider how the distinctive furrow pattern, which is thought to contribute to the final morphogenesis, is formed. We noticed that in the aforementioned analog simulation with paper, the distinctive pattern was created by folding the paper with varying shrinkage rates in vertical and horizontal directions. Therefore, we questioned whether the same process occurs in the actual phyllosoma-to-puerulus transformation. The initial morphological analysis and observations simplified the pre- and post-molt morphology, revealing differences in shrinkage rates in vertical and horizontal directions (Fig. S1). By approximating the pre-molt shape as an ellipse and the post-molt shape as a rectangle, we can model the transformation using a simple formula. The density distribution of the elliptical shape packed into the rectangle was considered, with the potential outcome of this configuration being the formation of the distinctive pattern. Initially, displacements of ellipses and rectangles were positioned on each side of the rectangle. For oblique sites that did not fit the rectangle, elements were placed at the vertices proportionally, based on vertical and horizontal excesses, and distributed along the edges following a Gaussian distribution. Subsequently, they were distributed inside the rectangle according to a Gaussian distribution in which the internal influence is varied by the variance parameter σ (see Fig. S2). The density gradient pattern of the distributed values varied depending on the parameter σ of the Gaussian distribution (Fig. 4a). When σ is small, the pattern stops on the porch, and when σ is large, it penetrates more inside. By adjusting the parameters, we could create a pattern similar to that seen in the actual molting process (Fig. 4a). Assuming that buckling occurs at a certain threshold when density changes, it is suggested that the furrow pattern during molt may be caused by differences in shrinkage rates. Furthermore, the toy model created this time allows for alterations to the ellipticity of the ellipse and aspect ratio of the rectangle. An investigation was conducted to ascertain the impact of varying these parameters on the final pattern. For the σ, the value was selected to reproduce the similar pattern to actual furrow observed during molting. As a consequence, it was determined that alterations to the ellipticity of the ellipse and aspect ratio of the rectangle resulted in corresponding changes to the final pattern (Fig. 4b, c). Of particular interest is the observation that as the disparity in shrinkage rates in vertical and horizontal directions decreases, the angle of the pattern becomes more gradual. In comparison to Palinuridae, the nisto larvae and adults of Scyllaridae have a slower curvature of the carapace and a more circular shape of the phyllosoma larvae (Landeira, Deville et al. 2023). This may be explained by our toy model, suggesting that further investigation into carapace changes in Scyllaridae phyllosoma larvae is warranted.

**Fig. 4.**
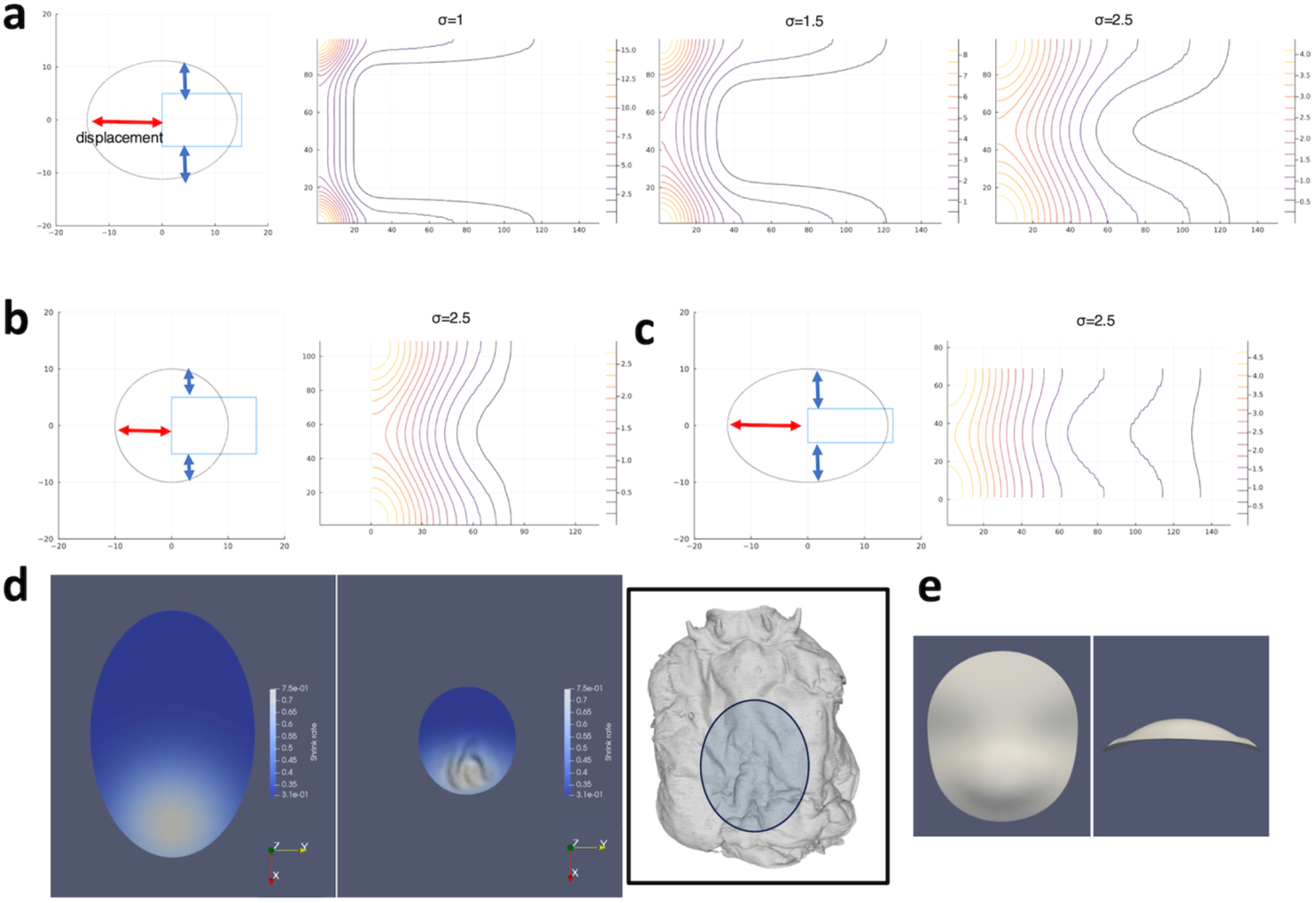
Reproducing furrow patterns in two and three dimensions. (a) Displacement concentration pattern produced when the dorsal carapace of phyllosoma and puerulus larvae is approximated as an ellipse and a rectangle, and the displacement is distributed inside the rectangle. (b) Displacement concentration pattern produced when the ellipticity is set to 1. (c) Displacement concentration pattern produced when the rectangle is made thinner. (d) Pre-shrinkage and post-shrinkage morphology from simulations, along with folding observed in actual dorsal carapace primordia. (e) Image of the unfolded morphology contracted by the simulation (center of (d)).

The aforementioned toy models focused solely on two-dimensional patterns. In practice, whether varying shrinkage rates at different locations allow folding in three dimensions remains unclear. A three-dimensional shrink model has been used to simulate the remodeling of beetle horns during pupal-adult transition (Matsuda, Adachi et al. 2024). Therefore, we employed a three-dimensional shrink model to examine whether folding occurs under anisotropic and gradient shrinkage, with greater contraction along the long axis. Anisotropic shrinkage was set so that the overall area decreased by a factor of 0.4, with an additional 2/3 reduction along the long axis compared to the short axis and the limitation of out-of-plane deformation was also imposed. The results demonstrated that ectopic contraction with constrained out-of-plane deformation can form similar folding as observed in the lobster metamorphosis process (Fig. 4d). Furthermore, within this model, we demonstrated that unfolding the fold (which corresponds to contraction without restricting out-of-plane deformation) can lead to curvature (Fig. 4e). These results suggest that folding could also occur in real phyllosomes as a result of contraction and restricted out-of-plane deformation. Current knowledge about the limitation of out-of-plane deformation is limited, but it may be influenced by apical Extracellular Matrices (ECMs), such as cuticles. On the other hands, the folding of the carapace was observed in individuals that died during molting, and caution is required regarding whether this reflects the actual developmental phenomenon. At least, it is evident that the disparity in longitudinal and transverse contraction rates gives rise to the formation of a curved structure. Furthermore, while the present study has described and examined the deformation of the surface structure, it cannot be denied that deformation within the internal structure exerts some influence upon the surface structure. Nevertheless, it remains a fact that deformation can be described by considering the surface structure alone. In this study, the measurement of the mechanical parameters to be incorporated into the model proved technically challenging. Conducting experiments on the mechanics of crustaceans (e.g. nanoindentation) may refine the model in the future.

## Conclusion

In this study, our focus was on the deformation of the carapace, from that of the phyllosoma larvae to that of the puerulus larvae. First, using three-dimensional micro-computed tomography on fixed tissue, we accurately identified the deformation, including that of the carapace. Subsequently, the dynamics occurring in the carapace during the actual molting process were elucidated through live observation. Furthermore, scanning electron microscopy (SEM) observation of the carapace and carapace primordia, in conjunction with digital-analog simulations, indicated that the distinctive furrow pattern on the cuticle surface may contribute to the final deformation. Furthermore, by approximating the deformation of the carapace from an ellipse to a rectangle, we demonstrated that a distinctive furrow pattern can be generated by the difference in the shrinkage rate in vertical and horizontal directions. Additionally, we found that various patterns emerge by deforming the ellipse and rectangle, suggesting potential applicability to other species. We also demonstrated that in three dimensions, local contractions can generate folds that curve upon unfolding. This study enhances our understanding of arthropod molting morphogenesis and offers new insights into the morphological evolution of Achelata.

## Acknowledgments

We appreciate Mr. Taisuke Takenouchi (Organization Mie Prefectural Science and Technology Promotion Center) for sampling phyllosoma and puerulus larvae. We also appreciate the member of Pattern formation laboratory (Osaka University) including Dr. Keisuke Matsuda for helpful supports and discussion. We also thank Prof. Shizue Ohsawa laboratory (Nagoya University) and G-language group (Institute for Advanced Biosciences) for helpful supports and discussion. This research was supported in part by MEXT KAKENHI Grant Number 19K22428 (to SK), 20H05941 (to SK) and by “Innovation inspired by Nature” Research Support Program, SEKISUI CHEMICAL CO. LTD (to SK) and Mishima Kaiun Memorial Foundation. HA was supported by research funds from the Yamagata Prefectural Government and Tsuruoka City, Japan.

## Author Contributions

Research design: H.A. and S.K., Funding acquisition: S.K., Formal analysis: H.A. and K.M., Methodology: H.A., K.M., Y.I., Writing—original draft: H.A., Writing—review and editing: all authors

## Data availability

Additional details required to reproduce or reanalyze the data reported in this study are available from the authors upon request.

---

Movie1: Flat structure of the phyllosoma of *Panulirus japonicus*. https://drive.google.com/file/d/1VJJu1Z4pD9e933MCt7-eu1PLysccvmmP/view?usp=drive_link

Movie2 : Entire molt process from phyllosoma to puerulus of *Panulirus japonicus*. https://drive.google.com/file/d/1cdP7yguSViCYV0bHS28trx4SpwGarv8g/view?usp=drive_link

Movie3 : Carapace contraction. https://drive.google.com/file/d/1H0f-dtDqXLo89XG3J4jJfrH7MtYjAi0n/view?usp=drive_link

Movie4 : Pumping movement. https://drive.google.com/file/d/1sC3g3lnTJe0C08wwD2H2K6ynxLx1_w0v/view?usp=drive_link

**Fig. S1.**
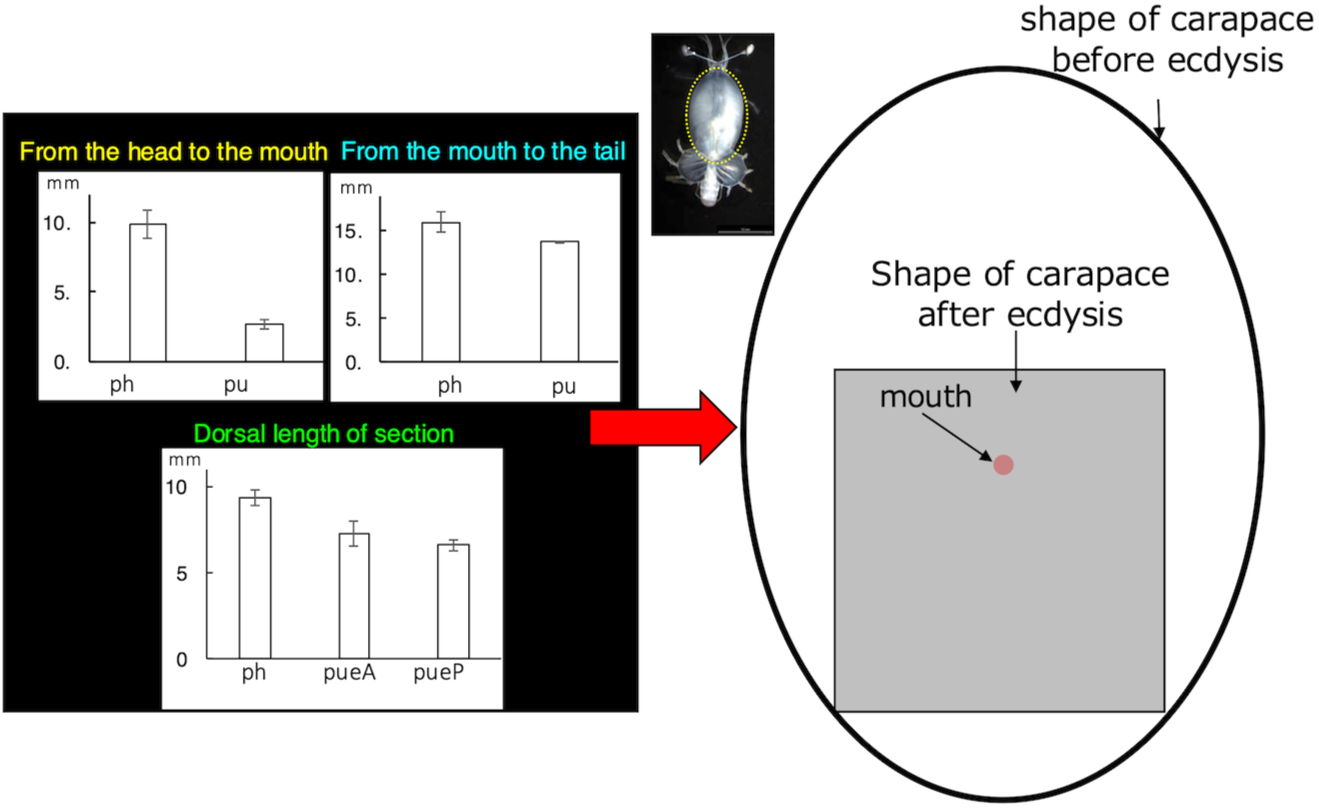
Elliptical and rectangular approximations of the carapace of phyllosoma and puerulus. The morphology of the phyllosoma and puerulus carapace was approximated as elliptical and rectangular, based on the morphological analysis in Fig. 1.

**Fig. S2.**
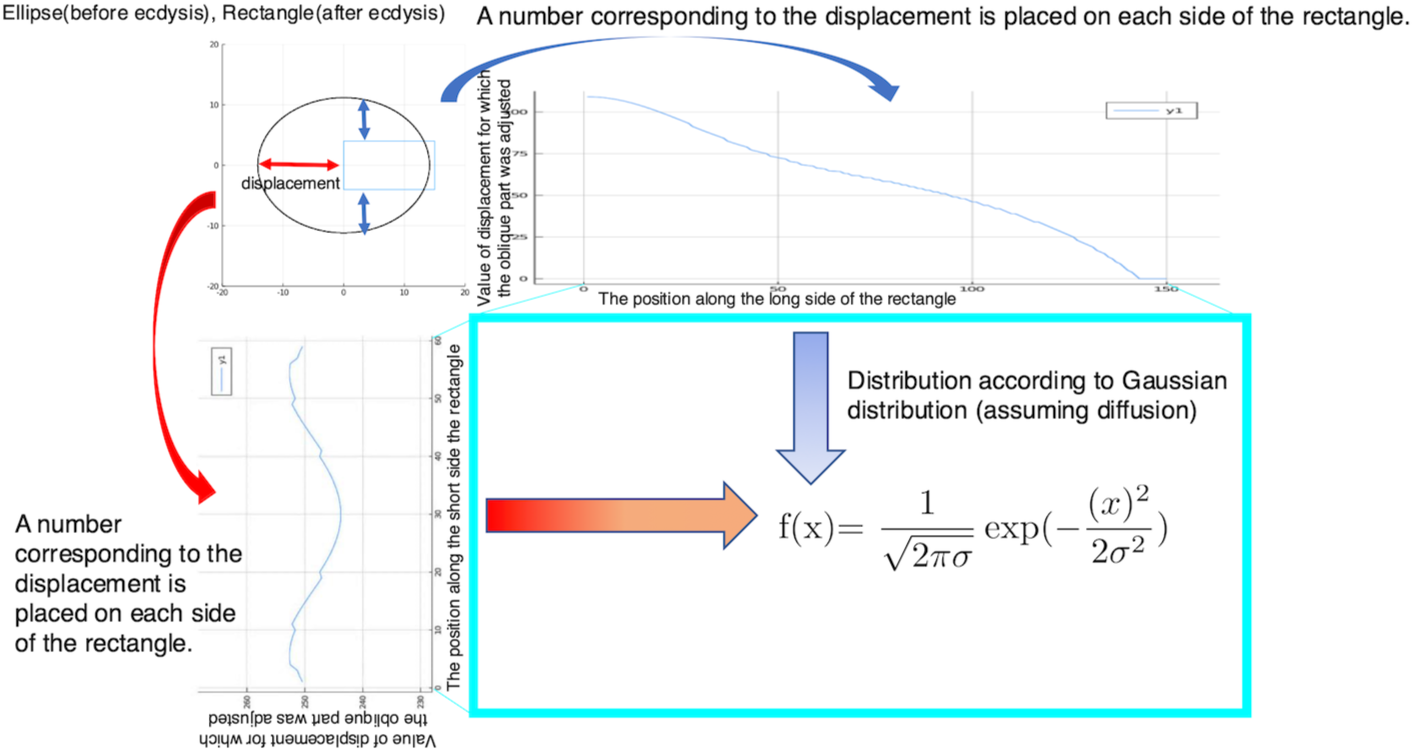
Methods of the contraction thought experiment in two dimensions. Rectangular and elliptical displacements were applied to each side of the rectangle and distributed within it according to a Gaussian distribution. For oblique sites that did not fit the rectangle, they were placed at the vertices proportionally, based on vertical and horizontal excesses, and distributed along the edges following a Gaussian distribution.

